# Transmission of prion infectivity from blood by peptoid-conjugated beads

**DOI:** 10.1101/609966

**Authors:** Simone Hornemann, Petra Schwarz, Elisabeth J. Rushing, Michael D. Connolly, Ronald N. Zuckermann, Alice Y. Yam, Adriano Aguzzi

## Abstract

Prions cause transmissible infectious diseases in humans and animals and have been found to be transmissible by blood transfusion even in the presymptomatic stage. However, the concentration of prions in body fluids such as blood and urine is extremely low, and therefore direct diagnostic tests on such specimens often yield false-negative results. Quantitative preanalytical prion enrichment may significantly improve the sensitivity of prion assays by concentrating trace amounts of prions from large volumes of body fluids. Here we show that beads conjugated to positively charged peptoids not only captured PrP aggregates from plasma of prion-infected hamsters, but also adsorbed prion infectivity in both the symptomatic and preclinical stages of the disease. Bead absorbed prion infectivity efficiently transmitted disease to transgenic indicator mice. We found that the readout of the peptoid-based misfolded protein assay (MPA) correlates closely with prion infectivity in vivo, thereby validating the MPA as a simple, quantitative, and sensitive surrogate indicator of the presence of prions. The reliable and sensitive detection of prions in plasma will enable a wide variety of applications in basic prion research and diagnostics.

## Introduction

Transmissible spongiform encephalopathies (TSEs) or prion diseases are caused by conformational transitioning of the physiological host-encoded prion protein, PrP^C^, into a misfolded and aggregated form, PrP^Sc^ that accumulates in the brain in the form of plaques or smaller plaque-like deposits [1, 2]. PrP^Sc^ differs from PrP^C^ in several biochemical and physicochemical properties such as its insolubility and partial resistance to proteinase K[1].

Prion diseases are prevalent in several animals and in humans. In humans, prion diseases can either appear spontaneously [3], be inherited [4] or acquired by infection [5–7] and include Creutzfeldt-Jakob disease, Gerstmann-Sträussler-Scheinker syndrome, fatal familial insomnia and Kuru. In the late 1990s a new form of CJD, designated variant CJD (vCJD) has emerged, which was traced to the transmission of prions to humans from cattle afflicted with BSE (bovine spongiform encephalopathy), indicating that BSE prions have overcome the species barrier between cattle and humans [8–11]. The possibility of the transmission of prions via blood was demonstrated by four confirmed fatal cases of vCJD in the UK that were acquired by blood transfusion [12, 13]. Moreover, studies performed in the rodents, sheep and deer, additionally showed that prions can be efficiently transmitted via whole blood transfusion as well as buffy coats from animals in the preclinical and clinical stages of the disease [14–16].

The concentration of prions in blood, however, is low. In hamsters, where it has been studied in detail, it reaches at most 10 LD_50_ units per ml blood (LD50 represents the lethal dose where 50% of the animals inoculated with prions come down) [17, 18]. This fact has made it difficult to utilize conventional transmission bioassays for detecting the presence of prion infectivity and has significantly hampered the validation of industrial procedures for prion removal from blood products. Assays that are based on *in vitro* amplification steps to increase the amount of detectable prions were developed [19, 20]. Efficient amplification and detection of PrP^Sc^ from the blood of hamsters infected with scrapie in the presymptomatic stage was demonstrated with the protein misfolding cyclic amplification (PMCA) assay [21]. More recently, the real-time quaking-induced conversion assay was developed, which amplifies PrP^Sc^ from tissue, cerebrospinal fluid (CSF), and other biological fluids [22, 23]. However, assays that are based on amplification steps are time-consuming and not well suited for automation and high-throughput screening applications. Also, assays that do not rely on the use of proteinase K (PK), which significantly reduces the total amount of aggregates available for detection, should allow for more sensitive detection of prions in body fluids such as blood.

An assay without additional amplification steps and PK digestion is the misfolded protein assay (MPA) [24–28]. The principle of the assay is based on a capturing step using magnetic peptoid-conjugated beads (PSR1) [24, 28, 29] that avidly and selectively bind to prion aggregates followed by quantification in an enzyme-linked immunosorbent assay (ELISA). The readouts of the MPA were found to correlate well with prion infectivity [30], suggesting that PSR1 beads physically capture prion seeds. Moreover, the MPA was able to detect PrP^Sc^, which was otherwise undetectable by proteinase K western blot analysis in the brain of a patient with familial CJD [25]. To further evaluate the reliability and sensitivity of the MPA for the detection of prion infectivity, we investigated in the present study the efficiency of PSR1 beads to capture prion infectivity from the plasma of hamsters that were in the presymptomatic and symptomatic stages of the disease. Additionally, we studied their efficiency to transmit disease to transgenic hamster indicator mice.

## Materials and Methods

### PSR1 beads

The PSR1 beads were generated by chemical conjugation of a thiolated PSR1 peptoid derivative to magnetic beads (Dynabeads™ M-270 Carboxylic Acid, Invitrogen, Carlsbad CA) as described previously and were provided as a 30mg/ml suspension of beads in bead storage buffer (1xPBS with 0.1% Sodium Azide, 0.01% Triton X-100) [28]. The PSR1 peptoid loading on the beads was measured by quantitative ninhydrin assay to be 9.88 nmol/mg of beads.

### Bead injections

12 μl or 120 μl of PSR1 beads (3 μl or 30 μl beads/mouse for a group of 4 mice) were incubated overnight with brain homogenate from healthy CD1 (wt) mice. Unbound material was removed by washing five times with 1 ml PBS. Beads were resuspended in 120 μl PBS and inoculated intracerebrally (i.c.) into *tga20* mice with groups of 3 mice (3 μl beads in a total volume of 30 μl/mouse) and 4 mice (30 μl beads/mouse). After 31 days post inoculation (dpi), mice were sacrificed and histopathologically examined for bead accumulation and toxicity.

### Preparation of hamster plasma

Golden Syrian hamsters at one-month weanling stage were inoculated intraperitoneally (i.p.) with 100 μl of 1% (w/v) 263K hamster prion strain infected hamster brain homogenate (estimated at 10^7^ LD_50_ infectious units) [31]. Control hamsters were inoculated with 100 μl of a 1% non-infectious hamster brain homogenate. Hamsters were sacrificed at the presymptomatic stage (30, 50 and 80 dpi) and at the symptomatic stage (104-106, 117-118, 143, 154 dpi). Control animals inoculated with PBS were sacrificed at 80 dpi. Blood was taken in the presence of EDTA-anticoagulant by cardiac puncture at various time points. Individual blood samples were centrifuged at 950xg for 10 minutes and the plasma in the supernatant fraction was transferred to another tube and frozen at −80°C. Hamsters were maintained under conventional conditions, and all experiments were performed at Novartis Diagnostics, Emeryville, USA in accordance with their animal welfare policies.

### Bead-based capture of PrP^Sc^

For the sensitivity assay, 30 μl of serial ten-fold dilutions of a 263K-infected 10% brain homogenate in PBS were inoculated i.c. into *Tg*(SHaPrP) mice (groups = 4-8) [32]. For the PSR1 capture assay, the plasma from 20 pre-symptomatic hamsters sacrificed at 30 dpi, from 20 presymptomatic hamsters at 50 dpi, from 14 symptomatic hamsters at 104-106 dpi, from 15 symptomatic hamsters at 117-118 dpi, from 11 symptomatic hamsters at 143 and 154 dpi, and from 2 non-infected hamsters at 80 days inoculated with PBS were combined to individual pools. 21 μl of PSR1 beads were washed five times in 1 ml PBS (8 mM Na_2_HPO_4_, 1.5 mM KH_2_PO_4_, 137 mM NaCl, 2.7 mM KCl, pH7.4) before incubation with 500 μl of pooled hamster plasma overnight at 4°C upon shaking. Beads were washed five times with 1 ml of PBS or TBSTT to remove unbound material and resuspended in 60 μl or 120 μl PBS, respectively. 30 μl of the resuspended beads were inoculated i.c. into *Tg*(SHaPrP) mice with groups of at least 4 mice.

For the analysis of the bead location in a *tga20* mouse inoculated with RML6-coated PSR1 beads, PSR 1 beads were treated with 10% RML6 brain homogenate ranging from 10^−2^ to 10^−10^ and i.c. inoculated into *tga20* mice overnight as previously described [30].

Mice were monitored every second day, and TSE was diagnosed according to clinical criteria including ataxia, wobbling, and hind leg paresis. At the onset of terminal disease Tg(SHaPrP) mice were sacrificed. Mice were maintained under conventional conditions, and all experiments were performed under license 130/2008 and in accordance to the regulations of the Veterinary Office of the Canton Zürich.

### Histopathology and immunohistochemical stains

Two-μm thick sections were cut onto positively charged silanized glass slides and stained with hematoxylin and eosin (HE) or immunostained using antibodies for PrP (SAF84), astrocytes (glial fibrillary acidic protein, GFAP) and microglia (IBA-1). For PrP staining, sections were deparaffinized and incubated for 6 min in 98% formic acid, then washed in distilled water for 5 min. Sections were heat-treated and immunohistochemically stained on an automated NEXES immunohistochemistry staining apparatus (Ventana Medical Systems, Switzerland) using an IVIEW DAB Detection Kit (Ventana). After incubation with protease 2 (Ventana) for 16 min, sections were incubated with anti-PrP SAF-84 (SPI bio; 1:200) for 32 min. Sections were counterstained with hematoxylin. GFAP immunohistochemistry for astrocytes (rabbit anti–mouse GFAP polyclonal antibody 1:13000 for 24 min; DAKO) and IBA-1 for microglia (rabbit polyclonal antibody 1:1000 for 24 min, WAKO) was similarly performed.

Histoblot analysis was performed using a modified standard protocol according to [33]. Briefly, 10 μm thick cryosections were mounted on glass slides and immediately pressed to a Nitrocellulose membrane (Protran, Schleicher & Schuell), soaked with lysis buffer (10 mM Tris, 100 mM NaCl, 0.05% Tween 20, pH 7.8) and air dried. After protein transfer, sections were rehydrated in TBST for 1 hour previous to Proteinase K digestion with 20, 50 and 100 μg/ml in 10 mM Tris-HCl pH 7.8 containing 100 mM NaCl and 0.1% Brij35), for 4 hours at 37 °C. After washing the membrane 3 times in TBST, a denaturation step with 3 M Guanidinium thiocyanate in 10 mM Tris-HCl, pH 7.8 was performed for 10 min at room temperature. The membrane was washed and blocked with 5% non-fat milk (in TBST) and incubated with anti-PrP antibody POM-1 (epitope in the globular domain, aa 121-231), 1:10000, overnight at 4 °C [34]. The blots were washed again and an alkaline-phosphate-conjugated goat anti mouse antibody was added (DAKO, 1:2000). Another washing step with TBST and B3 buffer (100 mM Tris, 100 mM NaCl, 100 mM MgCl_2_, pH 9.5) was followed by the visualisation step with BCIP/NBT (Roche) for 45 minutes. The colour development step was stopped with distilled water. Blots were air-dried and pictures were taken with an Olympus SZX12 Binocular and Olympus Camera.

### Immunoblotting

10% brain homogenates were prepared in 0.32 M sucrose by using a Precellys24 (Bertin). Extracts of 50-90 μg protein were digested with 50 μg/ml proteinase-K in DOC/NP-40 0.5% for 45 minutes at 37°C. The reaction was stopped by adding 3 μl complete protease inhibitor cocktail (7x concentrated) and 8 μl of a lauryl dodecyl sulfate (LDS)-based sample buffer. The samples were heated to 95°C for 5 minutes prior to electrophoresis through a 12% Bis-Tris precast gel (Invitrogen), followed by transfer to a nitrocellulose membrane by wet blotting. Proteins were detected with anti-PrP POM-1 antibody (1:10000). For secondary detection, an HRP-conjugated anti-mouse IgG antibody (Zymed, Invitrogen) was used and signals were visualized with an ECL detection kit (Pierce).

## Results

### Assessment of bead toxicity

In a pilot experiment, we first evaluated, whether peptoid-conjugated beads exposed to brain homogenates from healthy wild-type CD1 mice would induce any acute toxic response in recipient mice (Table 1). 3 μl or 30 μl beads, respectively, from a 30 mg/ml suspension of beads were incubated overnight with the brain homogenate and inoculated i.c. into *tga20* mice overexpressing murine PrP^C^ [35]. All mice were sacrificed at 31 days, except for one mouse that had already died at 8 dpi. All mice were analysed for acute toxicity and bead accumulation. In the brain of the mouse, that had died at 8 dpi, beads were detectable only at the injection site (Fig. 1A, S1 Figure), whereas in the remaining mice they were found to be broadly distributed in both hemispheres. Beads were often observed to be grouped in small clusters and were more abundant in intra- and periventricular regions, including the corpus callosum, ependymal lining, choroid plexus as well as the leptomeninges (Fig. 1B-E, S2-S3 Figure). Mild gliosis and microglial activation were observed around the injection site at 8 dpi (Fig. 1A), while a more diffuse astrogliosis was observed at 31 dpi (Fig. 1B-E, S2-S3 Figure). Gliosis, which is a nonspecific reaction to injury in the central nervous system [36], was more pronounced in the hippocampus, deep gray nuclei, medial habenular nucleus, brainstem and deep cerebellar white matter. As the lateral ventricles did not show evidence of hydrocephalus, we conclude that the beads did not significantly disturb the outflow of the cerebrospinal fluid (S1 Figure).

**Fig. 1.**
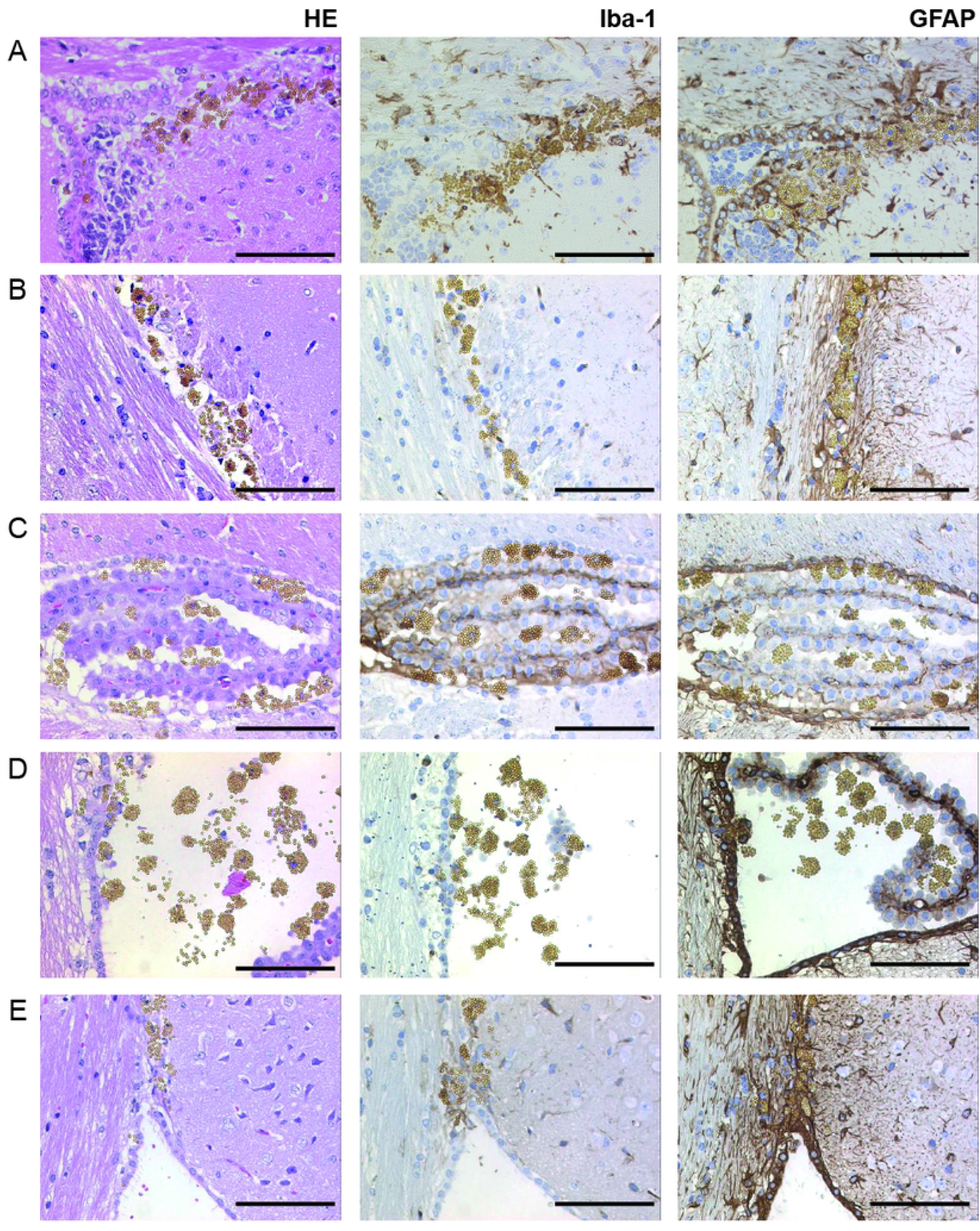
Histopathological analysis of tissue reactions to intracerebral bead inoculation. *Tga20* mice were inoculated with PSR1 beads that had been exposed to wt CD1 brain homogenates. Vacuolation by hematoxylin-eosin staining (HE), astrocytic gliosis was documented by an antibody directed against the glial fibrillary acidic protein (GFAP), and microgliosis was assessed with the activated microglial marker (Iba-1). **(A)** Brain sections of a *tga20* mice analysed at 8 dpi showed bead accumulation near the injection site. **(B)** Brain sections analyzed at 31 dpi showed accumulation of the beads adjacent to the corpus callosum, **(C)** choroid plexus and **(D)** lateral ventricle, and in the area between corpus callosum and the septofimbrial nucleus **(E)**. (Scale bars: 100 μm).

**Table 1:** Pilot experiment: Assessment of bead toxicity in *tga20*.

| PSR1 beads, coated with healthy brain homogenate | Total inoculated | Sacrificed at |
| --- | --- | --- |
| 3 µl beads in 30 µl PBS/mouse | 3 | 31, 31, 31 dpi |
| 30 µl in 30 µl PBS beads/mouse | 4 | 8*, 31, 31, 31 dpi |

\*Mouse died of an intercurrent death.

### End-point titration of the 263K hamster strain in *Tg*(SHaPrP)

10-fold serial dilutions (10^−2^ to 10^−12^) of a 10% (w/v) 263K hamster brain homogenate were i.c. inoculated into groups of 4 to 8 *Tg*(SHaPrP) mice, overexpressing the wild-type hamster prion protein [32] (Fig. 2A, Table 2). Clinical signs were observed in mice inoculated with dilutions spanning 10^−2^ to 10^−9^ after mean incubation periods between 40 and 107 days (Fig. 3, Table 2). From these data, a standard calibration curve for the infectivity of the 263K hamster strain in *Tg*(SHaPrP) was generated and a median 50% lethal dose [LD_50_] of 10^9.23^ units ml^−1^ using the Reed-Muench formula [37–39] was calculated.

**Fig. 2.**
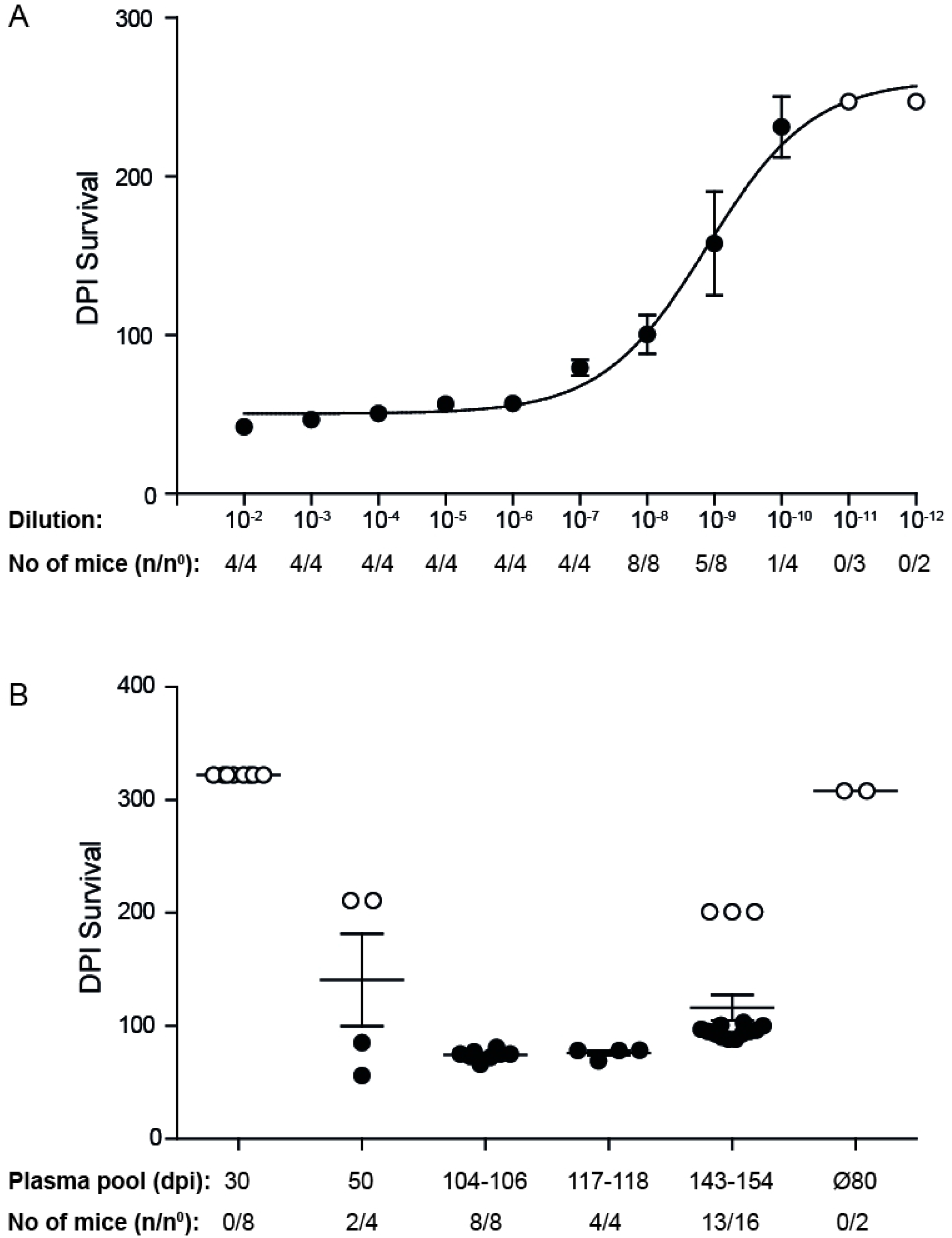
Survival plots of *tg*(SHaPrP) mice. **(A)** For titer determination, *tg*(SHaPrP) mice were inoculated with serial 10-fold dilutions of a 10% (wt/vol) 263K hamster brain homogenate ranging from 10^−2^ to 10^−12^. **(B)** Bioassay of *Tg*(SHaPrP) mice that were inoculated i.c. with PSR1 beads incubated with pooled plasma samples from hamsters infected with 263K prions. Individual pools of plasma samples from pre-symptomatic hamsters scarified at 30 (n=20) and 50 dpi (n=20) and from symptomatic hamsters scarified at 104-106 dpi (n=14), 117-118 dpi (n=15), and 143 and 154 dpi (n=11) were used. Data points: mean incubation times ± standard error of the mean. Mice that did not develop a prion disease until the end of the experiment are indicated by open circles. n/n^0^: indicates the attack rate (number of mice developing a prion disease divided by the total number of inoculated mice). Ø80: indicates the control group of mice inoculated with PSR1 beads after incubation with plasma from healthy 80 days old hamsters (n=2).

**Fig. 3.**
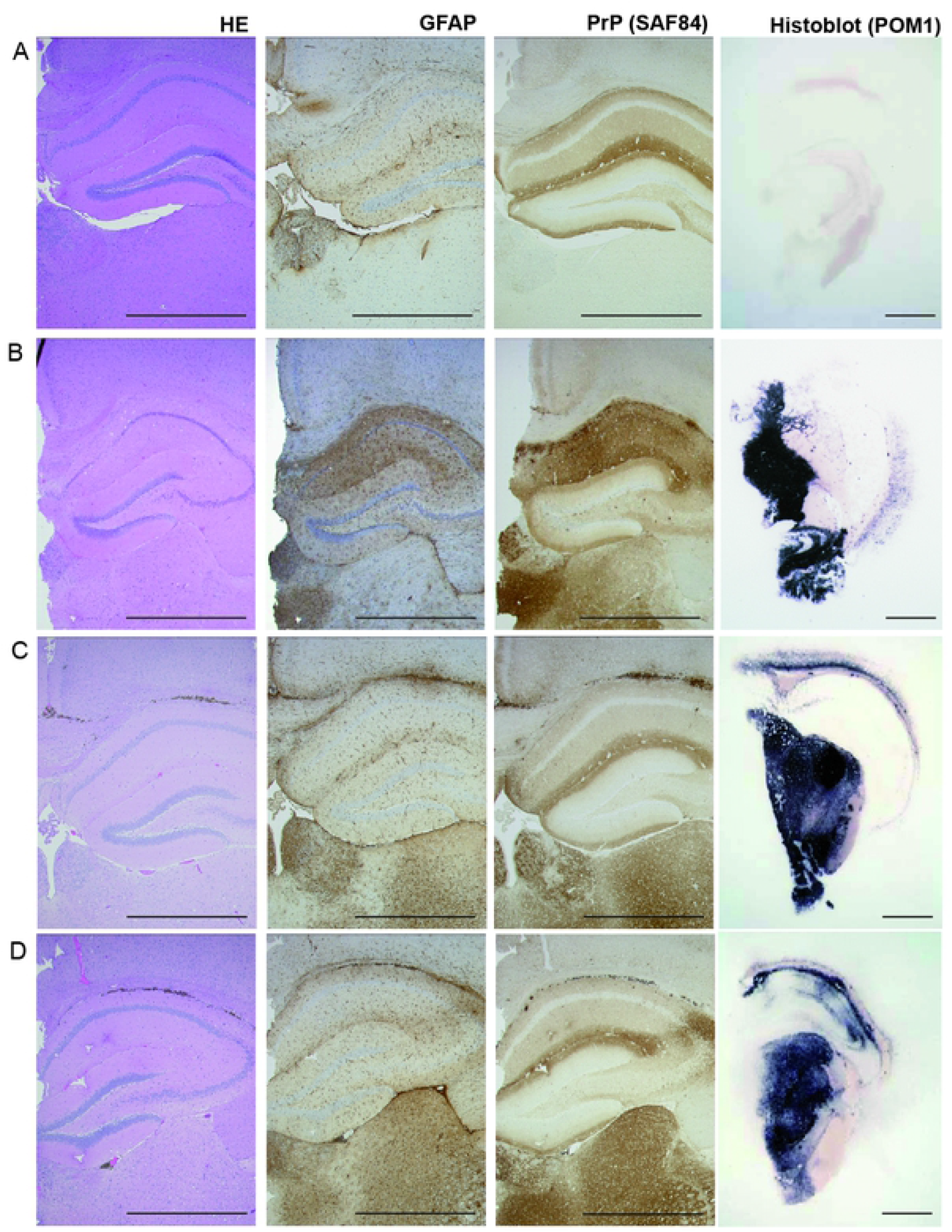
Pathology of hippocampus, medial habenular and thalamic nuclei from *Tg*(SHaPrP) mice. Non-inoculated mice **(A)** show no evidence of vacuolation, PrP^Sc^ deposition or gliosis. Mice inoculated with 263K prion-infected hamster brain homogenate **(B)**, inoculated with PSR1 beads incubated with plasma pools from symptomatic hamster at 117-118 dpi **(C)** and at 143 and 154 dpi **(D)** show vacuoles in the HE stained section (see also Figs. S2 and S3), PrP^c^ and PrP^Sc^ deposition as visualized by the PrP antibody SAF84 and astrocytic gliosis as evidenced by an antibody directed against GFAP. Histoblot analysis was used to show PrP^Sc^ deposition after proteinase K digestion and staining with POM1. (Scale bars: 1 mm).

**Table 2:**
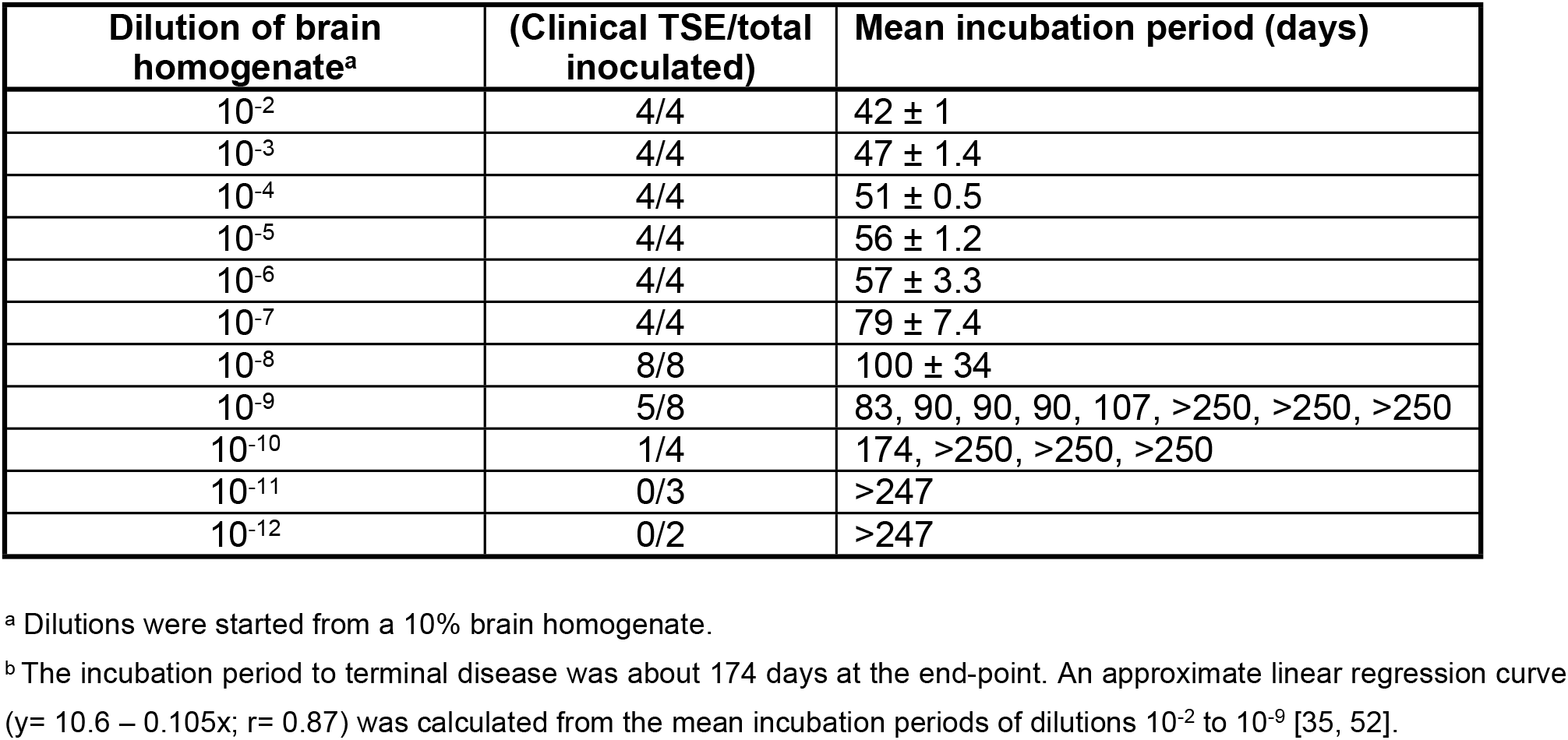
Summary of end-point titrations of the 263K inoculum in *Tg*(SHaPrP)

| Dilution of brain homogenate<sup>a</sup> | (Clinical TSE/total inoculated) | Mean incubation period (days) |
| --- | --- | --- |
| 10<sup>-2</sup> | 4/4 | 42 ± 1 |
| 10<sup>-3</sup> | 4/4 | 47 ± 1.4 |
| 10<sup>-4</sup> | 4/4 | 51 ± 0.5 |
| 10<sup>-5</sup> | 4/4 | 56 ± 1.2 |
| 10<sup>-6</sup> | 4/4 | 57 ± 3.3 |
| 10<sup>-7</sup> | 4/4 | 79 ± 7.4 |
| 10<sup>-8</sup> | 8/8 | 100 ± 34 |
| 10<sup>-9</sup> | 5/8 | 83, 90, 90, 90, 107, >250, >250, >250 |
| 10<sup>-10</sup> | 1/4 | 174, >250, >250, >250 |
| 10<sup>-11</sup> | 0/3 | >247 |
| 10<sup>-12</sup> | 0/2 | >247 |

<sup>a</sup> Dilutions were started from a 10% brain homogenate.
<sup>b</sup> The incubation period to terminal disease was about 174 days at the end-point. An approximate linear regression curve ( $y = 10.6 - 0.105x$ ; $r = 0.87$ ) was calculated from the mean incubation periods of dilutions 10<sup>-2</sup> to 10<sup>-9</sup> [35, 52].

### Bioassay in *Tg*(SHaPrP) mice of PSR1 beads treated with plasma from prion infected hamster

Next, we determined in a bioassay in Tg(SHaPrP) transgenic mice whether the PSR1 beads were able to capture quantitatively prion infectivity from plasma samples. To that purpose, we used pools of plasma from 263K prion-infected hamster at presymptomatic or symptomatic stages. Plasma samples from symptomatic hamsters were collected at the stage of onset of clinical signs with the typical disease characteristics of ataxia, loss of appetite and poor grooming. The individual pools (Table 3) were incubated overnight with the PSR1 beads. After an intense washing step, the beads were i.c. inoculated into Tg(SHaPrP) mice [32]. None of the mice inoculated with PSR1 beads coated with plasma from presymptomatic hamsters (30 dpi) nor any of the negative control mice showed any clinical signs of a prion disease. However, two mice (n= 4) inoculated with beads incubated with plasma from a presymptomatic hamster (50 dpi) developed disease with highly variable incubation periods after 56 and 85 dpi (equivalent to an apparent LD_50_ of 3.2 ± 2.2 units ml^−1^ of 30 μl 10% 263K hamster brain homogenate, Fig. 2B, Table 3) and incomplete attack rate.

Most other mice inoculated with beads treated with the different plasma pools from symptomatic hamster showed mean incubation periods between 74 ± 3 and 94 ± 5 dpi (equivalent to apparent LD_50_’s of 2.8 ± 0.3 to 1.9 ± 0.5 units ml^−1^ Fig. 2B, Table 3). The incubation times within the transgenic mouse bioassay increased with the use of inoculums obtained from symptomatic hamster with longer incubation times. The longest incubation time in the indicator mice was observed for the pooled inoculum derived from symptomatic hamster sacrificed at days 143 dpi and 154 dpi. This group of mice also exhibited a lower attack rate of 81%.

**Table 3:** Summary of bioassays with *Tg*(SHaPrP) mice that were inoculated with plasma coated PSR1 beads.

| **Plasma Pools** | **Attack rate**<br>**(Clinical TSE/total**<br>**inoculated)** | **Mean incubation period**<br>**(days)** | **Estimated titers\***<br>**Log LD<sub>50</sub> units ml<sup>-1</sup>** |
| --- | --- | --- | --- |
| pre-symptomatic<br>hamster, 20 animals,<br>30 dpi | 0% (0/8) | 8 survivor terminated at 322<br>dpi |  |
| pre-symptomatic<br>hamster, 20 animals,<br>50 dpi | 50% (2/4) | 56 and 85 dpi (71 ± 21 dpi)<br>(plus 2 survivor terminated<br>at 211 dpi) | 3.2 ± 2.2 |
| symptomatic<br>hamster, 14 animals,<br>104-106 dpi | 100% (8/8) | 74 ± 3 dpi | 2.8 ± 0.3 |
| symptomatic<br>hamster, 15 animals,<br>117-118 dpi | 100% (4/4) | 76 ± 5 dpi | 2.4 ± 0.5 |
| symptomatic<br>hamster, 11 animals,<br>143 dpi and 154 dpi | 81% (13/16) | 94 ± 5 dpi<br>(plus 3 survivor terminated<br>at 211 dpi) | 1.9 ± 0.5 |
| non-infected<br>hamster, 2 animals,<br>80 days after PBS<br>inoculation | 0% (0/2) | 8 survivor terminated at 308<br>dpi |  |

\*Titers were estimated from the calibration curve of the end-point titrations of the 263K inoculum in *Tg*(SHaPrP) (Table 2; Fig. 2) using the incubation time method [35, 52].

The calculated titers are slightly higher than the apparent titer of 1.25 LD_50_ ml^−1^, estimated from the quantity of 10 LD_50_ ml^−1^ infectivity found in the blood of hamsters during the symptomatic stage of the disease [17, 40], for 30 μl resuspended PSR1 beads coated with plasma from prion-infected hamster. Histopathological and immunohistochemical analysis of the brains of the mice was used to diagnostically confirm the presence for a prion disease (Fig. 3). Additionally, PK resistant PrP, a surrogate marker of prion disease, could be detected by histoblot and Western blot analyses (Fig. 3 and 4). We therefore conclude that PSR1 beads highly efficiently capture prion infectivity from plasma from presymptomatic and symptomatic cases and are able to transmit infectivity to *Tg*(SHaPrP) mice.

**Fig. 4.**
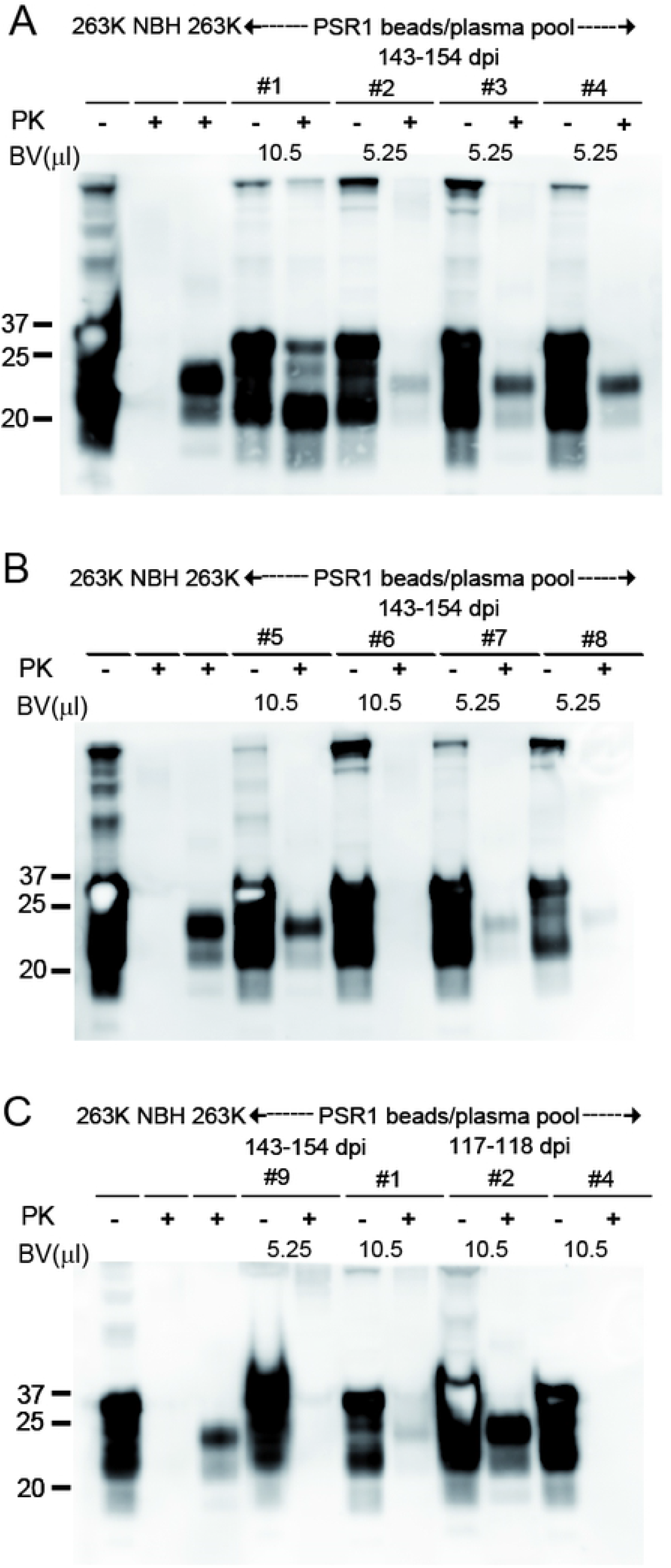
PK-Western blot analysis of brain homogenates from *Tg*(SHaPrP) mice inoculated with PSR1 beads coated with plasma from symptomatic hamster. **(A-C)** PrP^Sc^ is present in *Tg*(SHaPrP) inoculated i.c. with PSR1 beads incubated with plasma pools from symptomatic hamsters. Mice were incubated either with 5.25 or 10.5 μl of PSR1 beads in PBS (A) or in TBSTT (B-C). Because of the high expression level of PrP^C^ in *Tg*(SHaPrP), PrP^C^ is still detectable on Western blots. Control samples are labeled with NBH: non-infected brain homogenate from healthy mice, and 263K: brain homogenate from mice inoculated with 263K prions. BV: bead volume. The anti-PrP antibody POM-1 was used for detection. The molecular weight standard is shown in kilodaltons.

Mice were also analyzed after termination for the location of the beads. After 90 dpi, beads were mostly found in the brain, spinal cord and CSF, but also in most other organs (if only in small numbers) as well as in urine and feces (see S1-3 Table). After 250 dpi, only a few beads were observed in the brain with occasional beads around the spinal cord, in the spleen and in selected other organs (see S1-3 Table). A similar distribution of the beads was also observed in *tga20* mice i.c. inoculated with PSR1 beads treated with a 10^−9^ dilution of a 10% (w/v) RML6 brain homogenate and sacrificed at 253 dpi (S4 Figure, [30]). These data indicate that beads might have been transported along the axonal route [41] or lymphatics [42, 43].

## Discussion and Conclusion

Previously, we have reported that the MPA could detect prion aggregates with high sensitivity in the brain of a patient with a familial prion disease [25]. Crucially, the MPA detected PK sensitive conformers of PrP^Sc^ that were only barely detectable by standard methods [25]. These findings prompted us to investigate further the sensitivity of the peptoid-conjugated beads to capture prion infectivity from the plasma of prion-infected hamsters. This is biologically and medically relevant because (1) the conformation of infectious PrP in blood is unknown and (2) the detection of prions in blood and blood products continues to represent a major analytical challenge.

Our data showed that the PSR1 beads were able to enrich low concentrations of infectivity from the plasma of hamsters in symptomatic and even presymptomatic stages of the disease. These findings indicate that the beads are capable of capturing prion infectivity from plasma of less than 10 LD_50_ ml^−1^, the quantity of infectivity identified in the blood of hamsters during the symptomatic stage [17, 40]. Prions in plasma were proposed to consist of more PK sensitive conformations of PrP^Sc^ [44], suggesting that the high capture efficiency of PSR1 beads could stem from an ability to detect more PK sensitive conformations of PrP^Sc^. Alternatively, the apparent higher titers of PrP^Sc^ on the bead surfaces might occur from PrP^Sc^ being protected against degradation on the beads.

Transmission of infectivity from solid-state surfaces might occur either by dissociation of infectivity or by the transport of these surfaces to different brain areas where they initiate infection [45, 46]. We observed that the beads stayed at the injection site within the first days after injection. After 31 days, the beads were found to be more diffusely distributed within the brain, with a greater density in periventricular regions and within the leptomeninges. After 90 days, only a few beads could be detected, which were subsequently found in most organs and body fluids. Possible routes of clearance include the internal and external veins that enter the dural sinuses, and then eventually drain into the internal jugular veins. Other potential conduits for clearance are along axons [41] or recently described lymphatic vasculature that lines dural sinuses in the meninges and drains into cervical lymph nodes [42, 43, 47].

The ability of PSR1-coated beads to efficiently capture and enrich prions from plasma directly suggests the feasibility of solid-state prion-capture matrices for both clinical and research uses. For example, the beads might be applied for depletion, removal and clearance of prions from biological fluids and biopharmaceutical products. The number of beads can easily be increased and their magnetic properties should allow the capture of prions even in high volumes.

In a further embodiment, the beads might also be applied for the preanalytical enrichment of prions in concert with the scrapie cell endpoint assay (SCEPA) [48] and/or with prion amplification assays [49]. Prion-concentrating steps in combination with these assays were recently reported to improve their sensitivity by several orders of magnitude [48, 50, 51].

Finally, the beads might also be used together with a sensitive detection method to diagnose prions. Prion bioassays report the infectious titer and do not directly reflect the total number of PrP molecules, whereas the MPA reports the amount of PrP in aggregated conformations. Because each aggregate can be composed of a vast number of PrP molecules, the subsequent disaggregation can lead to the exposure of many epitopes for sensitive immunodetection. A sensitive prion detection assay based on the MPA technology could be easily automatable and optimized for high-throughput applications, eventually paving the way to early and sensitive blood diagnosis of prion diseases in humans and to the prevention of transmission of prions through economical and reliable screening of blood and blood products.

## Abbreviations

PrP: prion protein
PrP^C^: cellular isoform of PrP
PrP^Sc^: scrapie isoform of PrP^C^
mPrP: mouse PrP
TSE: transmissible spongiform encephalopathy
PSR1: Prion Specific Reagent 1
RML6: Rocky Mountain Laboratory strain, passage 6
MPA: Misfolded Protein Assay
ELISA: Enzyme-Linked Immunosorbent Assay

## Acknowledgements

The authors thank Marianne König for technical assistance. AA is the recipient of an Advanced Grant of the European Research Council, and is supported by the Swiss National Foundation, the Clinical Research Priority Programs “Small RNAs” and “Human Hemato-Lymphatic Diseases”, SystemsX.ch, and the GELU Foundation. SH is the recipient of grants from SystemsX.ch (SynucleiX) and the innovations commission of the University Hospital of Zurich. Work at the Molecular Foundry was supported by the Office of Science, Office of Basic Energy Sciences, of the U.S. Department of Energy under Contract No. DE-AC02-05CH11231. This study was supported by the Novartis Research Foundation.

## Author Contributions

**Conceptualization:** AA SH.

**Formal analysis:** AA SH PS EJR.

**Funding acquisition:** AA.

**Investigation:** PS.

**Project administration:** AA SH.

**Resources:** MDC RNZ AYY.

**Supervision:** AA SH.

**Validation:** PS.

**Visualization:** AA SH PS.

**Writing ± original draft:** AA SH EJR.

**Writing ± review & editing:** All authors.

